# Prefrontal cortical activity predicts the extra-place field spiking of hippocampal place cells

**DOI:** 10.1101/2020.11.23.395012

**Authors:** Jai Y. Yu, Loren M. Frank

## Abstract

The receptive field of a neuron describes the regions of a stimulus space where the neuron is consistently active. Sparse spiking outside of the receptive field is often considered to be noise, rather than a reflection of information processing. Whether this characterization is accurate remains unclear. We therefore contrasted the sparse, temporally isolated spiking of hippocampal CA1 place cells to the consistent, temporally adjacent spiking seen within their spatial receptive fields (“place fields”). We found that isolated spikes, which occur during locomotion, are more strongly phase coupled to hippocampal theta oscillations than adjacent spikes and, surprisingly, transiently express coherent representations of non-local spatial representations. Further, prefrontal cortical activity is coordinated with, and can predict the occurrence of future isolated spiking events. Rather than local noise within the hippocampus, sparse, isolated place cell spiking reflects a coordinated cortical-hippocampal process consistent with the generation of non-local scenario representations during active navigation.

## Main text

The concept of a receptive field (Sherrington 1906, Hartline 1938, Spillmann 2014) provides a fundamental model for how neural spiking can convey information about features in the external environment. In the hippocampus, many cells show spatially tuned receptive fields (O’Keefe and Dostrovsky 1971, O’Keefe 1976). The spiking rate of these “place cells” rises and then falls as an animal traverses specific locations in an environment. Locations with high spiking rates are defined as a cell’s “place fields” (O’Keefe 1976, McNaughton, Barnes et al. 1983), and place field-associated spiking of place cells conveys sufficient spatial information to estimate the animal’s location with high accuracy (Muller and Kubie 1989, Wilson and McNaughton 1993, Brown, Frank et al. 1998, Zhang, Ginzburg et al. 1998).

Although the majority of place cell spiking occurs when an animal is moving within the cell’s place fields, occasional spiking occurs when the animal is at locations outside the fields (O’Keefe 1976, McNaughton, Barnes et al. 1983, Muller, Kubie et al. 1987, Thompson and Best 1989). These “isolated” spiking events can occur during movement and are distinct from sparse spiking observed during Sharp Wave/Ripples (SWRs) seen during immobility (Buzsáki 2015). Importantly, isolated spikes are not locked to specific locations. As a result, standard analyses that average activity across many passes through the same location (Olton, Branch et al. 1978, Hill and Best 1981, Thompson and Best 1989, Wilson and McNaughton 1993, Frank, Brown et al. 2000, Jensen and Lisman 2000) effectively exclude these spikes from further consideration. Whether these spikes reflect unreliable, noisy processes that merit exclusion or whether they instead reflect coherent, meaningful signals (Stein, Gossen et al. 2005, Faisal, Selen et al. 2008, Masquelier 2013) remains unknown.

Noise in neural networks can arise from stochastic cellular events that cause the membrane voltage to occasionally exceed the action potential threshold, even without upstream input (Stein, Gossen et al. 2005, Faisal, Selen et al. 2008). While the spatially selective inputs to a place cell raise the membrane voltage closer to the action potential threshold when an animal approaches the cell’s place field (Epsztein, Brecht et al. 2011), stochastic events causing occasional increases in membrane potential could result in spiking outside of a cell’s place field. However, previous observations indicate that at least some spiking outside of a cell’s typical place fields reflect mnemonic processes rather than noise. CA1 and CA3 place cells can emit spikes outside of their place fields as an animal approaches choice points (Johnson and Redish 2007, Kay, Chung et al. 2020) and during vicarious trial and error (Johnson and Redish 2007), or when an animal is travelling in the opposite direction over a location with a place field (Kay, Chung et al. 2020). These events are hypothesized to reflect non-current scenarios, such as simulating possible future scenarios when a decision needs to be made (Johnson, Fenton et al. 2009, Kay, Chung et al. 2020).

How can we determine whether isolated spiking outside of a place cell’s spatially tuned receptive field reflects information processing in the hippocampal circuit as opposed to activity that does not reflect information processing or noise? Spiking due to stochastic cellular events is expected to be local to individual neurons. By contrast, spiking associated with information processing would be expected to co-vary in a consistent manner across neurons in both local and distributed networks (Masquelier 2013). Thus, if spiking outside of the classical place field conveys information, we would expect it to (1) be coordinated across multiple hippocampal neurons, (2) contain coherent spatial information and (3) be coordinated with activity outside the hippocampus.

We therefore examined spiking both within the hippocampus and across the hippocampus and prefrontal cortex (PFC), focusing on activity during movement. The PFC is anatomically connected to the hippocampus through both direct and indirect projections (Swanson and Cowan 1977, Swanson 1981, Jay, Glowinski et al. 1989), and coordinated activity across these networks reflects their engagement during memory processing (Schacter, Addis et al. 2007, Preston and Eichenbaum 2013, Eichenbaum 2017). For example, network level coherence between prefrontal cortex and hippocampus increases during periods when memory retrieval occurs (Hyman, Zilli et al. 2005, Jones and Wilson 2005, Benchenane, Peyrache et al. 2010, Sigurdsson, Stark et al. 2010, Place, Farovik et al. 2016, Guise and Shapiro 2017, Myroshnychenko, Seamans et al. 2017, Zielinski, Shin et al. 2019). Whether PFC activity differs systematically at the time of isolated spiking in the hippocampus remains unknown.

Our examination of isolated spiking of place cells revealed that these events reflect the coherent activation of hippocampal representations of physically distant locations, and that these events are coordinated with ongoing activity in the PFC. These findings suggest that isolated spikes are a signature of distributed and coherent information processing in the brain.

## Results

In order to understand the extent of isolated spiking during active behavior and to identify a potential function of this activity, we took an unbiased approach where we surveyed CA1 place cell spiking across all movement periods (animal speed >2cm/s) as animals performed a spatial navigation task (Yu, Kay et al. 2017, Yu, Liu et al. 2018) (Figure 1A-B, Figure S1). In the hippocampus the temporal structure of spiking during locomotion is strongly influenced by the endogenous ~8Hz theta rhythm (O’Keefe and Recce 1993), and bouts of higher rate spiking corresponding to place field traversals spanned multiple, adjacent cycles of theta (Fig. 1C). As expected, we also observed isolated spikes where a neuron would be silent for many theta cycles, emit a small number of spikes on a single theta cycle, and then return to being silent (Fig. 1D) (Olton, Branch et al. 1978, Hill and Best 1981, Thompson and Best 1989, Johnson and Redish 2007).

**Figure 1.**
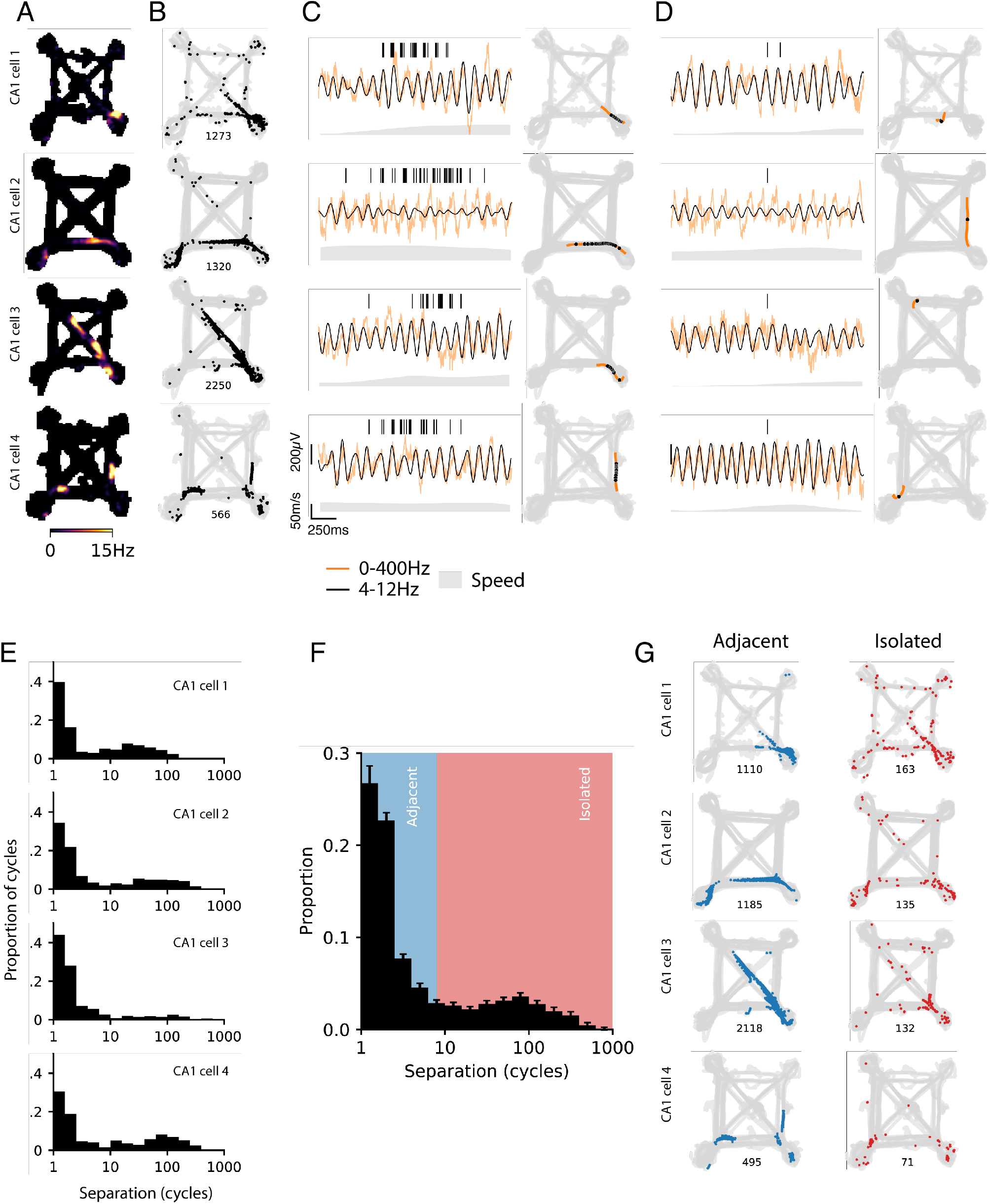
Isolated and adjacent spiking activity of hippocampal CA1 place cells. A. Occupancy normalized spiking rate maps for spiking activity during active movement (animal speed >2cm/s) across behavior sessions for each day for four example CA1 cells. B. Location of spiking (black dots) and animal trajectory (gray) for occupancy maps in A. Spike count shown below each panel. C. Spike raster and corresponding location for a bout of spiking activity over adjacent theta cycles. Raw (orange) and theta frequency filtered (black) local field potential are shown below the spike raster. The corresponding location on the maze for the bout is shown on the right. D. Spike raster and corresponding location for spiking isolated from other spiking activity. The corresponding location on the maze for the bout is shown on the right. E. Distribution of mean separation between theta cycles with spiking. Separation is defined as the mean cycle count to 3 nearest neighbor cycles with spiking. F. Population distribution of mean separation between a theta cycles (n=301 cells). G. Location of spiking classified as adjacent activity (<8 mean cycles of separation) or isolated activity (≥ mean 8 cycles of separation) for the 4 example cells in A. Spike count shown below each panel.

The standard approach to defining place field spiking relies on averaging spiking rates across many traversals of a location. This average provides a useful experimental summary of spiking, but information averaged across traversals is not directly available to downstream neurons. We therefore we used a criterion to distinguish between “adjacent” and “isolated” spiking based on the local temporal organization of spiking. Specifically, given the importance of theta in organizing hippocampal activity (Vanderwolf 1969, O’Keefe and Recce 1993, Buzsaki 2002), we calculated the interval between neighboring theta cycles with spiking (in cycles, mean of nearest three) (Fig. 1E, Fig. S2). As expected, the majority of spike-containing theta cycles are near another spike-containing cycle. The remaining spike-containing theta cycles are separated from neighboring spike-containing cycles by up to hundreds of cycles, reflecting their temporal isolation. When plotted on a log scale, the underlying distribution was bimodal, and based on this distribution we chose a threshold of 8 cycles of mean separation to each theta cycle with spiking to define “adjacent” or “isolated” activity (n=301 cells, Fig. 1F). This separation captured intuitive notions of within- and extra-place field activity: adjacent activity was spatially concentrated, as expected from place-field spiking (Fig. 1E); isolated activity was spatially sparse and lack the high spiking rates observed for place field activity (Fig. 1G). We also verified that isolated activity spikes, although sparsely emitted, were very unlikely to be spike clustering errors (Fig. S3).

One previously described form of extra-field spiking occurred around choice points and coincides with overt deliberation behavior (Johnson and Redish 2007). Isolated activity, in contrast, was not more frequent around choice point locations (Fig. S4A), nor were there differences in the speed (Fig. S4B) or angular velocity (Fig. S4C) of the animal at times of isolated as compared to adjacent spiking. Thus, isolated spiking is not restricted to specific active behavioral states or locations, such as path choice points. Isolated spiking was also not associated with SWRs, which are transient network oscillations observed in the local field potential (LFP) predominantly found when the animal is moving slowly or is immobile (Buzsáki 2015). We excluded SWRs from our analyses (see methods) and also independently confirmed the isolated spiking events did not occur during SWRs by comparing the hippocampal LFP associated with excluded spiking (low speed periods and SWRs) with the LFP associated with isolated spiking. The LFP associated with excluded spiking showed a network spectral signature consistent with SWRs (Fig. S5A left column, B), with power in the slow gamma (~30Hz) and ripple frequencies (~150-250Hz). In contrast, the LFP associated with isolated spiking shows a different network spectral signature, with power in the theta band (Buzsaki, Leung et al. 1983) (Fig. S5A center column, B). Indeed, the network spectral signature of isolated spiking is very similarity to the LFP associated with adjacent spiking, and even has slightly higher theta power (Fig. S5A right column, B).

Isolated spiking was also highly concentrated within each theta cycle, a potential signature of an information containing signal (Masquelier 2013). Place field-associated spiking displays strong phase-coupling to the hippocampal theta rhythm, where the maximum probability of spiking occurs in earlier phases near the trough of theta (Buzsaki, Leung et al. 1983, O’Keefe and Recce 1993). By contrast, the entrainment of isolated spiking preferentially occurred in the late phases of theta (Fig. 2A-B). Isolated spiking was also more tightly phase locked to theta compared with adjacent spiking (Fig. 2C).

**Figure 2.**
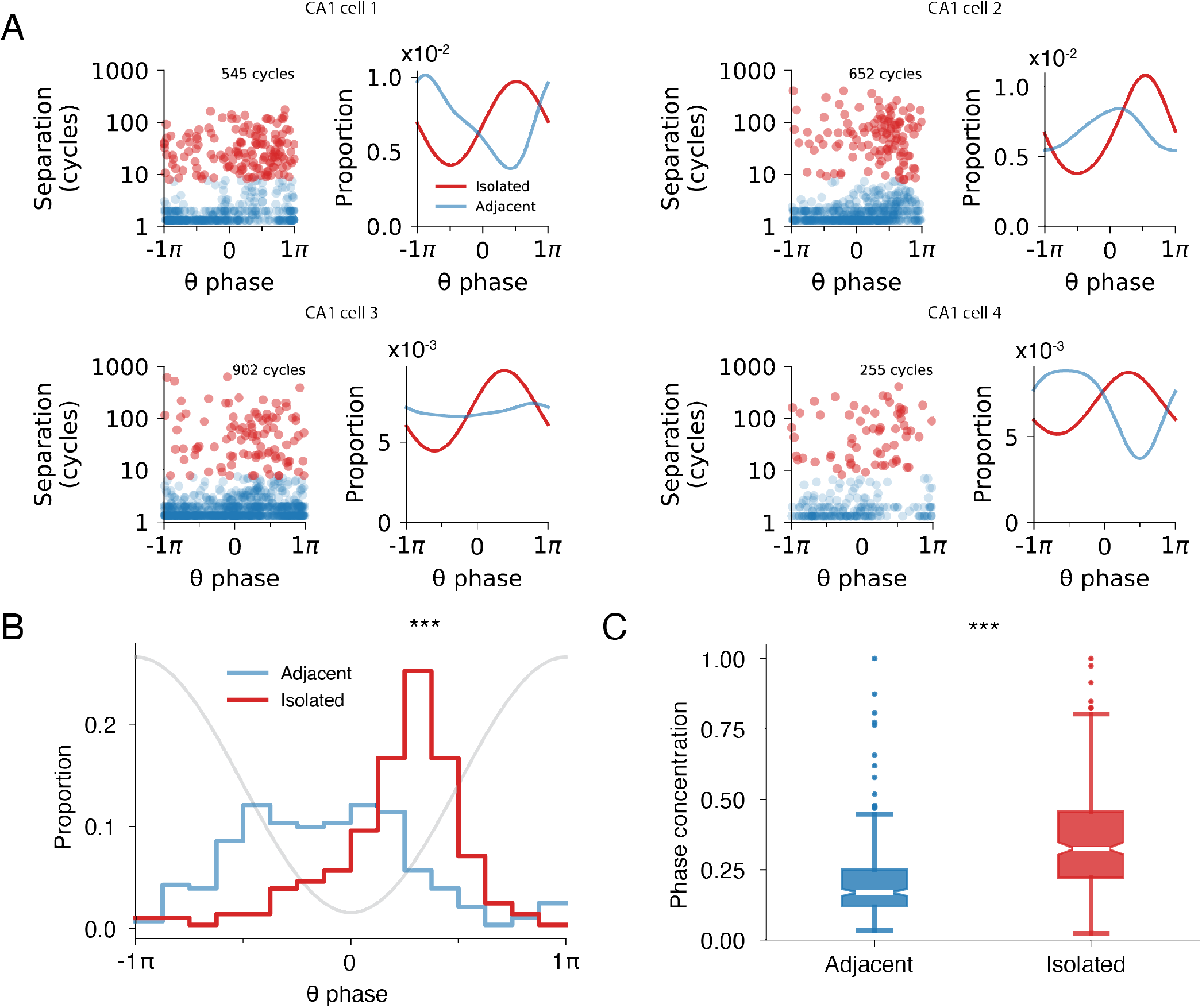
Isolated and adjacent spiking activity show distinct phase locking to hippocampal theta oscillation. A. Separation versus mean spike theta phase preference for each theta cycle with spiking. A separation threshold of 8 cycles between isolated and adjacent is based on Fig. 1. Histogram shows the mean spiking phase for each theta cycle. Examples correspond to the 4 cells from Fig. 1. Circular median test between the isolated and adjacent distributions: top left p=1.4×10^−5^; top right p=8.9×10^−8^; bottom left p=2.4×10^−4^; bottom right p=5.3×10^−2^. B. Mean theta phase preference distribution for adjacent and isolated spiking for the CA1 place cell population (n=279 cells). Gray line illustrates theta phase. Circular median test: p=0. C. Mean theta phase concentration distribution for adjacent and isolated spiking for the CA1 place cell population (n=279 cells). Wilcoxon rank-sum test: p=4.4×10^−26^

Spiking during the late phases of theta has been associated with the expression of non-local representations, including to be visited locations or locations previously visited (Skaggs, McNaughton et al. 1996, Redish 2016, Kay, Chung et al. 2020). Could isolated spiking be a signature of a coherent non-local representation? If so, then we would expect that pairs of neurons that are co-active during periods of adjacent spiking (e.g. cells likely to have overlapping place fields) would also be co-active within a theta cycle containing isolated spiking events. We examined this possibility by using an approach that has been used to demonstrate reactivation of non-local spatial representations during SWRs, where a pair of place cells is more likely to spike together if their place fields overlap (Karlsson and Frank 2009, Singer and Frank 2009) (Fig. 3A). First, we calculate the likelihood of co-spiking for a pair of place cells that had isolated spiking within the same theta cycle. We then quantified the overlap in their adjacent spiking activity.

**Figure 3.**
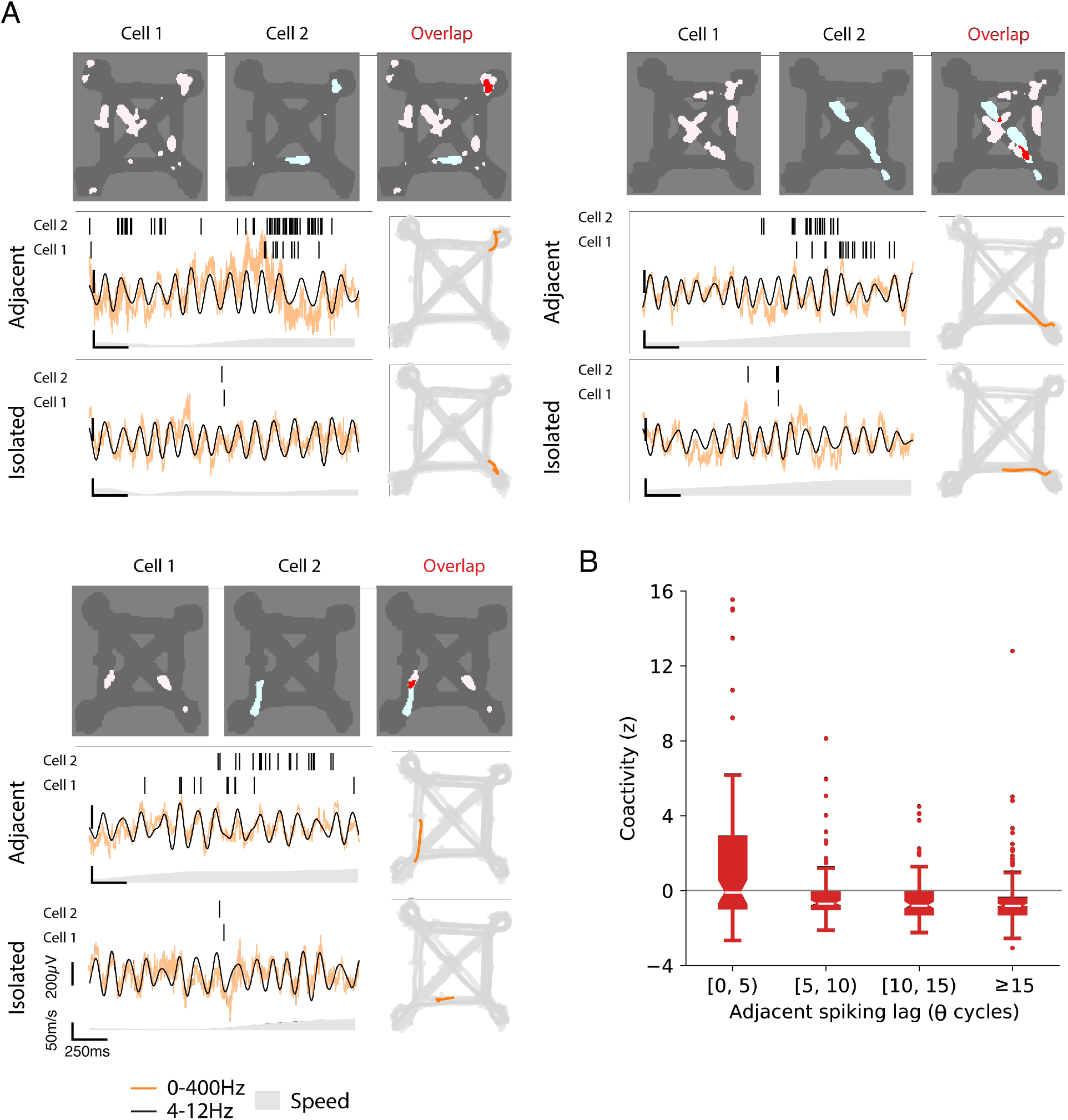
Reactivation of spatiotemporal place field activity relationships during theta cycles with isolated spiking activity. A. Three pairs of CA1 cells with overlapping adjacent activity. The place fields (occupancy normalize spiking rate > 5Hz) for each cell as well as their spatial overlap are shown. Example spiking bouts of adjacent and isolated activity are shown with raw and theta frequency band filtered LFP as well as their corresponding locations. B. Normalized coactivity (z) for CA1 cell pairs during theta cycles with isolated activity (n=425 pairs) grouped by the mean separation in time (mean lag) between their adjacent activity (R=-0.28, R^2^=0.077, p=6.40×10^−9^).

We found that cells that fired together during periods of adjacent spiking were also more likely to fire together during isolated spiking events. Across the population, lower lags in spiking during adjacent activity were correlated with greater co-spiking during isolated events (Fig. 3B, R=-0.28, R^2^=0.077, p=6.40×10^−9^). These findings support the notion that isolated spikes reflect the structured and coherent activation of a remote spatial representation.

As a further test of the hypothesis that isolated spiking contains information, we asked whether these spiking events in the hippocampus marked the time of specific activity patterns in PFC. Given the anatomical and functional connectivity between these regions, evidence of coordination between hippocampus and PFC would argue that these events are not the result of local noise in the hippocampus but instead reflect coherent and structured activity across brain regions. An example of such hippocampal-cortical engagement occurs during SWRs, where hippocampal reactivation is accompanied by the coordinated reactivation of cortical representations (Remondes and Wilson 2015, Jadhav, Rothschild et al. 2016, Wang and Ikemoto 2016, Rothschild, Eban et al. 2017, Yu, Kay et al. 2017, Yu, Liu et al. 2018). If such coordination is seen around the times of isolated spikes, we should be able to identify PFC neurons that spike differently around times of isolated activity in the hippocampus than at comparable periods where isolated spiking was not observed.

We first selected theta cycles with isolated spiking for a given CA1 cell. Next, we found matching theta cycles from other times when the animal was moving through the same locations in the same direction at a similar speed, but where the CA1 cell was not active (e.g. did not have isolated spiking) (Fig. S6). This was possible because, in our task, the animal traversed the a given location multiple times, providing a pool of theta cycles, of which only a subset contained isolated spiking. Importantly, none of the matching cycles contained adjacent spiking, confirming that the isolated spiking events were not simply events on the edge of a place field. We then compared the spiking of simultaneously recorded PFC neurons between cycles with isolated activity and these matched control cycles (Fig. 4A). We note that theta coordinates activity in hippocampal-cortical networks (Hyman, Zilli et al. 2005, Jones and Wilson 2005), allowing us to continue to use theta cycles as the temporal reference to relate activity across structures.

**Figure 4.**
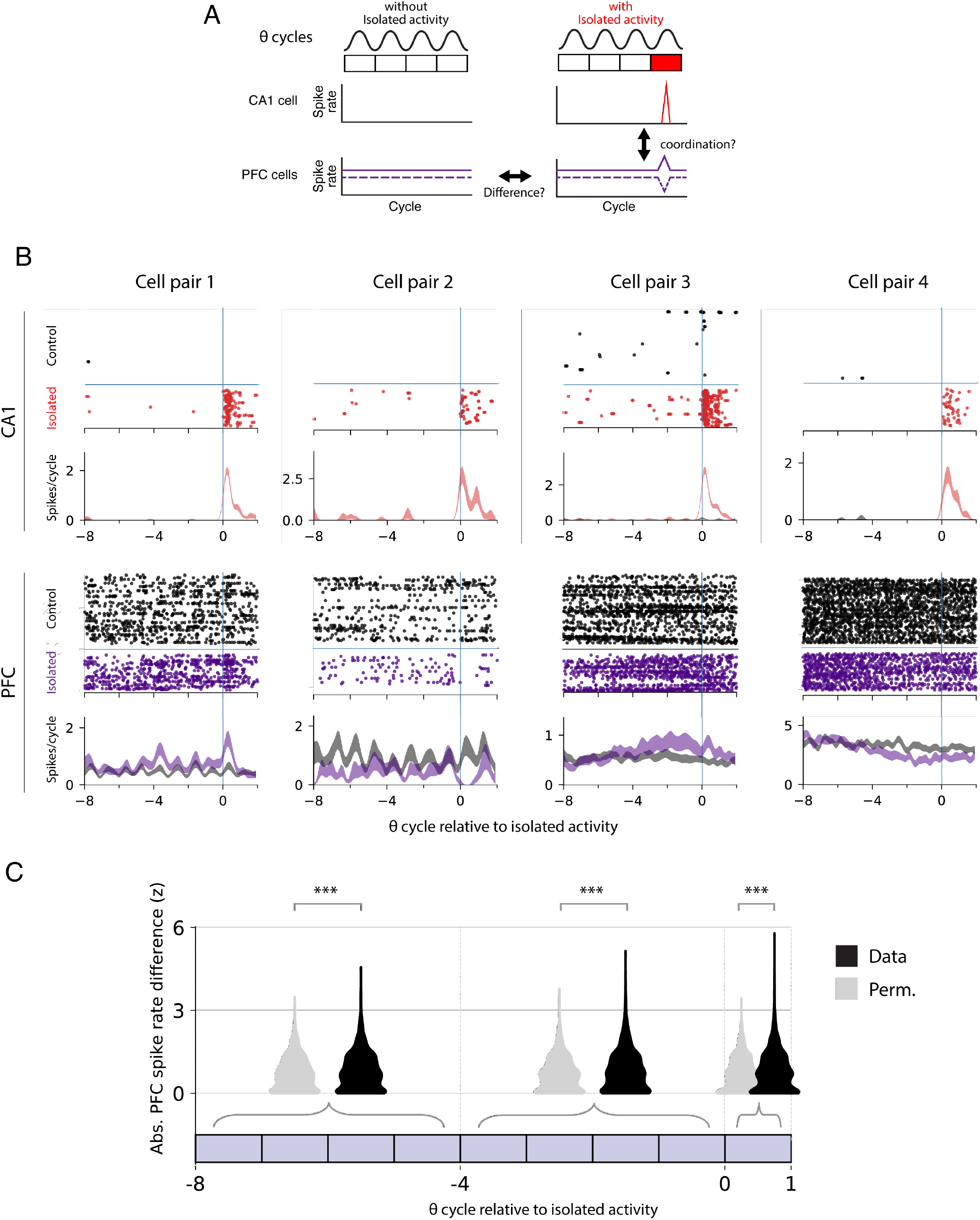
PFC activity is coordinated with hippocampal isolated activity. A. Schematic illustrating potential CA1 and PFC activity around the time of isolated spiking. Changes in PFC spiking around the time of CA1 isolated spiking may reflect coordination between the two regions. B. Example spike raster and spiking rate (Mean ± SEM) for pairs of co-recorded hippocampus CA1 and PFC cells. Each raster shows spiking aligned to isolated hippocampal activity (cycle 0) and matched control trials. Spiking is plotted relative to the cycles of the hippocampal theta rhythm. For CA1 cells, red indicates spikes and spiking rate for intervals with isolated spiking at cycle 0. Black indicates control intervals without isolated spiking at cycle 0. For PFC cells, purple indicates spikes and spiking rate for intervals with isolated spiking at cycle 0. Black indicates control intervals without isolated spiking at cycle 0. C. Violin plots and quantification of spike rate differences between control and actual intervals for PFC-CA1 cell pairs (n=2798) in time windows relative to CA1 isolated activity. Rate difference, original data (black) and permuted (gray), is expressed the z-score of the absolute observed difference relative to its own permuted distribution. The Wilcoxon signed-rank test (*** p<0.001) was used to compare the original and permuted groups: p=4.7×10^−8^, 3.6×10^−12^ and 2.2×10^−10^ for each group respectively.

We found PFC cells whose spiking rate differed depending on whether or not there was an associated period of isolated spiking for a given CA1 cell (Fig. 4B). We expect that only a small fraction of PFC cells would show a significant difference in spiking relative to the isolated spiking of a given CA1 cell, but nonetheless, across the population (n=2798 PFC-CA1 cell pairs), the difference in PFC firing rates between isolated and matched control periods was significantly larger than the permutation control (Fig. 4C). This difference indicates coordination between CA1 and prefrontal cortex around the time of CA1 isolated activity. Interestingly, this coordination was not limited to the specific isolated theta cycle: the difference remained significant even in a window of 4-8 theta cycles before the isolated spiking event, indicating that PFC activity could play a causal role in driving isolated spiking events in the hippocampus.

Under this hypothesis, the ensemble activity of PFC neurons should predict the future occurrence of hippocampal isolated activity (Fig. 5A). To test that prediction for a given CA1 cell, we used the spiking activity from all simultaneously recorded PFC ensembles (Median n=20, IQR=8 PFC cells per CA1 cell) to build cross-validated Generalized Linear Models with elastic net regularization. We compared the ability of the models to predict the occurrence of isolated activity related to a permutation control (see Methods, Fig. S7). We then carried out that analysis for all CA1 cells (n=158) with isolated spiking.

**Figure 5.**
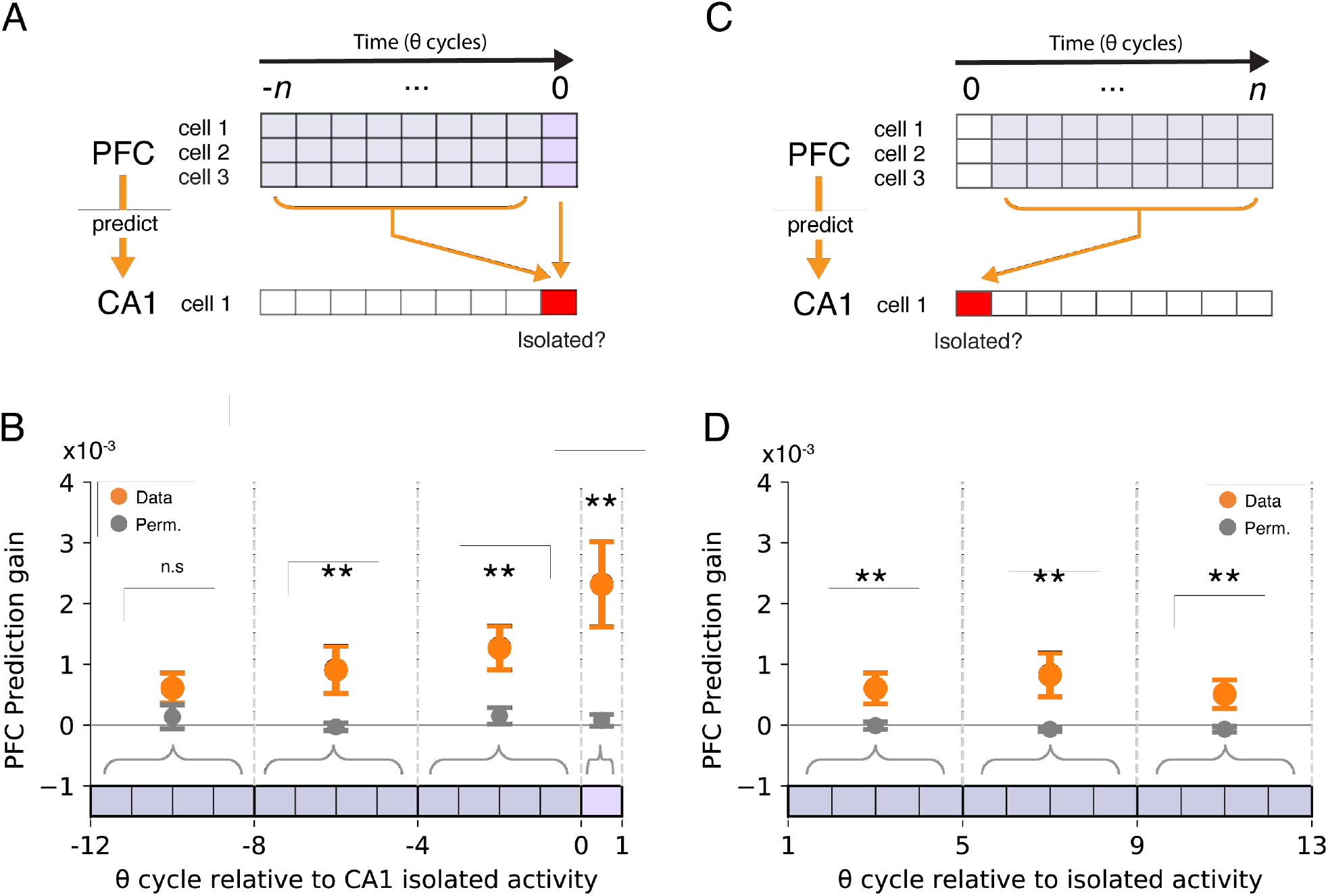
A. PFC activity leading up to isolated spiking is used to predict the future occurrence of isolated spiking in one CA1 place cell. B. Prediction gain (Mean ± SEM) of GLMs where PFC spiking activity is used to predict whether an upcoming CA1 theta cycle contains isolated or non-isolated control spiking activity for a given CA1 place cell (n=158). Pairwise permutation test (** p<0.05) for mean with multiple comparison correction: p=0.079, p=0.0027, p=0.0006 and p<0.0002 for each group respectively. C. PFC activity after isolated spiking is used to predict the previous occurrence of isolated spiking in one CA1 place cell. D. Prediction gain (Mean ± SEM) of GLMs where PFC spiking activity is used to predict whether a previous CA1 theta cycle contained isolated or non-isolated control spiking activity for a given CA1 place cell (n=162). Pairwise permutation test (** p<0.05) for mean with multiple comparison correction: p=0.0071, p=0.0002 and p=0.0053 for each group respectively.

We found that PFC activity can predict the occurrence of isolated spiking in CA1 at above chance levels, even in a window 4-8 theta cycles before the isolated spiking (Fig. 5B). We also asked whether there was any evidence consistent with isolated spiking in CA1 influencing subsequent PFC activity (Fig. 5C). We found that the coordination between hippocampus and prefrontal cortex persists after the occurrence of isolated activity but is weaker compared to intervals immediately before and during cycles with isolated activity (Fig. 5D). We also noted the average prediction gains were small in magnitude, which has previously been observed for prediction gains relating auditory and hippocampal activity around the times of SWRs (Rothschild, Eban et al. 2017). This is not surprising given the relatively small numbers of simultaneously recorded PFC cells that were available to predict the activity of any given CA1 unit. We can therefore regard these cross-validated predictions as lower bounds on the actual values that would be obtained if it were possible to sample the entire PFC population. Indeed, examining the values for individual PFC ensemble - CA1 models revealed a number of cases with prediction gains between 2.5 and 5% (Fig. S7). Thus, our results demonstrate that information expressed by prefrontal and hippocampal cell populations is coordinated around the time of isolated activity.

Importantly, the predictive PFC activity patterns were specific for individual CA1 cells. We examined the correlation between β-coefficients of PFC predictors across predictive models. If the spiking of specific PFC cells was strongly predictive of isolated spiking of a particular CA1 cell but not of other CA1 cells, this β-coefficient correlation should be low, indicating that a given PFC cell would predict the spiking in one model (e.g. one CA1 cell) but not another. By contrast, if a subset of PFC cells consistently predicted isolated spiking across CA1 cells, then these correlations would be high, as the same PFC cells would show similarly β-coefficients across models.

We found that the mean correlation coefficient was not significantly different from 0 (Median=-0.021, IQR=0.16, Wilcoxon rank sum test p=0.431). This indicates that the PFC ensembles predicting the occurrence of isolated activity for different CA1 cells are distinct, and argues for specificity in PFC-CA1 coordination around the occurrence of isolated activity.

## Discussion

We examined spiking outside of a place cell’s place field relative to local hippocampal network activity and to activity in prefrontal cortex. We found that this isolated spiking preferentially occurs during the late phase of theta oscillations, recapitulates coherent spatial representations, and is coordinated with prefrontal cortical activity. Our findings argue that seemingly spontaneous and sparse activity, previously considered as noise in the hippocampus, are actually precisely timed spikes that reflect coordinated activity both within the hippocampus and across the hippocampal-prefrontal networks.

We found evidence for CA1 isolated spiking reflecting structured information processing rather than noise within the hippocampal network at both the single cell and pairwise level of analysis. At the single cell level, isolated CA1 spiking is highly concentrated during in the late phases of theta, pointing to the segregation of current versus non-current scenarios between the early and late phase of theta respectively (Sanders, Renno-Costa et al. 2015). This is in line with previously described place cell spiking associated with non-local representations including possible future locations, travel in the non-current direction (Wang, Foster et al. 2020), and activity on non-preferred trajectories (Kay, Chung et al. 2020), all of which are seen preferentially during the late phases of theta. Our results also extend previous findings of non-local spiking associated with vicarious trial and error behavior seen near choice points or at the edges of place fields (Johnson and Redish 2007, Redish 2016). We found that isolated spikes occur throughout the environment and are not concentrated at the edges of place fields or near choice points. These spikes also occurred in association with high movement speeds. Pairwise analyses further demonstrated that isolated spikes are coordinated across hippocampal neurons: cells that fired together during adjacent spiking periods were also more likely to fire together within an isolated spiking event. This is consistent with a brief, coherent activation of a remote spatial representation, indicating that these events could support deliberative processes associated with the evaluation of distant physical locations.

Our analysis of isolated CA1 place cell spiking relative to PFC activity provided additional evidence that these events could support deliberative processes. At the single pair level, we identified individual PFC neurons that spiked differently in association with an isolated CA1 spiking event as compared to periods matched for location, direction of movement, and speed. At the ensemble level we found that these differences were significant not only during the theta cycle associated with isolated CA1 spiking, but also 4-8 cycles before the isolated spiking event. Moreover, ensemble PFC activity could predict the occurrence of a theta cycle with isolated CA1 spiking, and these predictions remained significant for PFC activity occurring 4-8 cycles before the isolated spiking. These predictions were also specific: a particular set of PFC cells were strong predictors of a given CA1’s cells isolated spiking, while a different set of PFC cells might predict isolated spiking in a different CA1 cell.

These findings indicate that isolated spiking events in CA1 are very unlikely to be the product of local, stochastic fluctuations in the hippocampus. Instead, our findings indicate the presence of a slow (~1s) change in PFC activity that could trigger an isolated spiking event in CA1. Interestingly, the changes in rate in PFC and the strength of the PFC-CA1 coupling is greatest during the cycle with isolated spiking but remains greater than expected by chance well after the CA1 spiking, suggesting the possibility that the CA1 spiking drives a subsequent change in PFC activity. Thus, our findings point to a potential cortical-hippocampal-cortical of information flow, conceptually similar to the cortical-hippocampal-cortical information flow seen around SWRs during sleep (Rothschild, Eban et al. 2017).

Our results also suggest that information exchange between cortex and hippocampus may occur frequently during active behavior. This extends previous findings of hippocampal-prefrontal coupling from imaging (Squire, Ojemann et al. 1992, Buckner, Petersen et al. 1995, Schacter, Alpert et al. 1996, Polyn, Natu et al. 2005, St Jacques, Kragel et al. 2011, Rugg and Vilberg 2013, Schedlbauer and Ekstrom 2019) and neural recording experiments (Tomita, Ohbayashi et al. 1999, Kyd and Bilkey 2003, Hyman, Zilli et al. 2005, Jones and Wilson 2005, Benchenane, Peyrache et al. 2010, Sigurdsson, Stark et al. 2010, Hok, Chah et al. 2013, Place, Farovik et al. 2016, Guise and Shapiro 2017, Myroshnychenko, Seamans et al. 2017, Zielinski, Shin et al. 2019) on the role of prefrontal cortex in modulating both cortical and subcortical structures during mnemonic processes (Tomita, Ohbayashi et al. 1999, Simons and Spiers 2003, St Jacques, Kragel et al. 2011, Eichenbaum 2017). Our results also complement findings demonstrating coherent spiking activity patterns across hippocampus and PFC in the context of both SWRs and locomotion-associated spiking (Jadhav, Rothschild et al. 2016, Shin, Tang et al. 2019, Zielinski, Shin et al. 2019).

Interestingly, the communication latency between PFC and HP is hypothesized to be ~26-28ms, or approximately ¼ of a theta cycle (Place, Farovik et al. 2016). We were therefore surprised to find PFC spiking can predict whether isolated spiking will occur up to 4-8 theta cycles, or approximately 500ms – 1s, later, an interval much longer than what is needed for direct information transfer. Although the channel for communication between PFC and hippocampus may have a short latency, our results suggest the expression of isolated hippocampal spiking likely involves coordinated activity between these regions (Squire, Ojemann et al. 1992, Buckner, Petersen et al. 1995, Schacter, Alpert et al. 1996) that evolves over time (Polyn, Natu et al. 2005, Rugg and Vilberg 2013, Schedlbauer and Ekstrom 2019). This is consistent with human imaging studies that show cortical activity change can precede memory recall on the order of seconds (Polyn, Natu et al. 2005). This long duration may in part be explained by the timescale of cortical processing where spiking time constants are >100ms (Murray, Bernacchia et al. 2014). The long timescale may reflect additional intracortical communication necessary to integrate information across multiple theta cycles, which eventually triggers the expression of hippocampal representations.

We hypothesize that these exchanges may serve to modulate ongoing hippocampal-cortical network representations corresponding to current experience with internally generated representations corresponding to non-current scenarios. Additionally, we hypothesize that through this mode of communication, PFC could drive the expression of non-current scenario representations from memory in the hippocampus, which in turn, feeds back to cortical regions as a part of an evaluation loop (Yu and Frank 2015). This coordination could be involved in covert evaluation of potential trajectories or goal locations for decision making in the future. The cortical drive could potentially underly previously reported extra-place field spiking and non-current spatial representations in the hippocampus associated with approach to a choice point (Kay, Chung et al. 2020), during vicarious trial and error (Johnson and Redish 2007), and spiking during travel in the non-preferred direction of place fields (Kay, Chung et al. 2020).

Both cortical and hippocampal spiking patterns display noise-like variation, even when the animal performs repeated tasks or actions (Tolhurst, Movshon et al. 1983, Fenton, 1998 #2585, Shadlen and Newsome 1998). However, cortical discharge can also be highly reproducible given a consistent input (Mainen and Sejnowski 1995), and behaviors can reflect a degree of accuracy consistent with very low levels of noise in the brain (Osborne, Lisberger et al. 2005). In the context of signal versus noise, our findings indicate that the sparse spiking of hippocampal place cells is better understood as a signal that express non-current representations consistent with alternative scenarios. These transient injections of non-current representations could signal processes that update ongoing hippocampal representations. Thus, the hippocampal place cell spiking during active behavior may dynamically reflect both externally driven and internally generated, non-current representations, which we hypothesize can collectively guide ongoing behavior.

## Author contributions

J.Y.Y and L.M.F. designed the analysis and wrote the manuscript. J.Y.Y analyzed the data.

## Acknowledgements

We thank A. Comrie, A. Gillespie, J. Guidera, A. Joshi and K. H. Lee for comments on the manuscript, and D. Liu, A. Loback and I. Grossrubatscher for assisting in the collection of the original data. This work was supported by a Jane Coffin Childs Memorial Fund for Biomedical Research postdoctoral fellowship (J.Y.Y.), the Howard Hughes Medical Institute, and University of California Office of the President Lab Fees Award #LF-12-237680 (L.M.F.).

## Data Availability

Data used for this manuscript can be accessed at: https://crcns.org/data-sets/hc/hc-13/about-hc-13

## Code Availability

Statistical analysis was performed using standard MATLAB, Scipy, Numpy modules listed in the methods and text. Figures were generated using MATLAB and Matplotlib. The code is available upon request.

---

**Figure S1.**
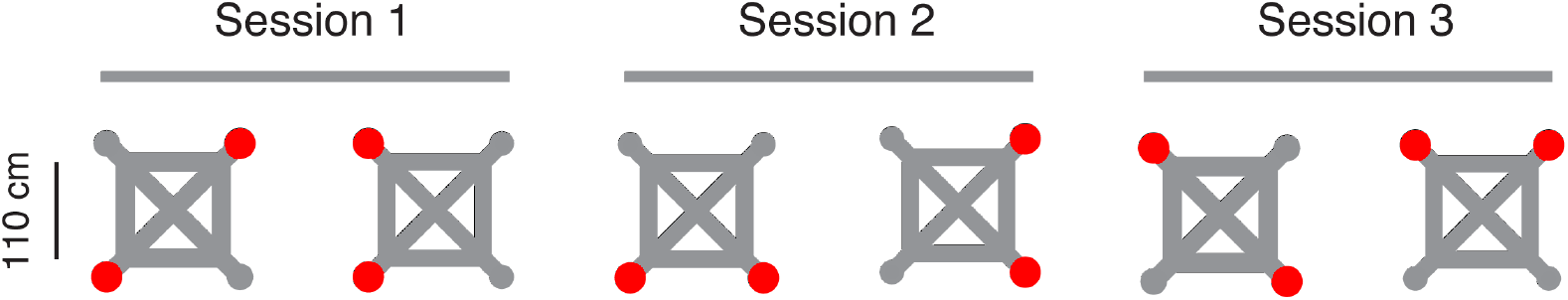
Task schematic. The maze has 4 potential reward locations (wells) located on the corners of the maze but only 2 will deliver reward for a given contingency. The animal needs to find the 2 wells that deliver reward (red circles) and alternate between the two to continuously receive reward. All reward wells are connected by paths (gray) and the rat is free to choose any path. The locations of the rewarded wells change within session, between sessions and/or between days. The pairs of wells that are rewarded vary across sessions and the rat needs to learn the locations of the rewarded wells for every contingency by trial and error.

**Figure S2.**
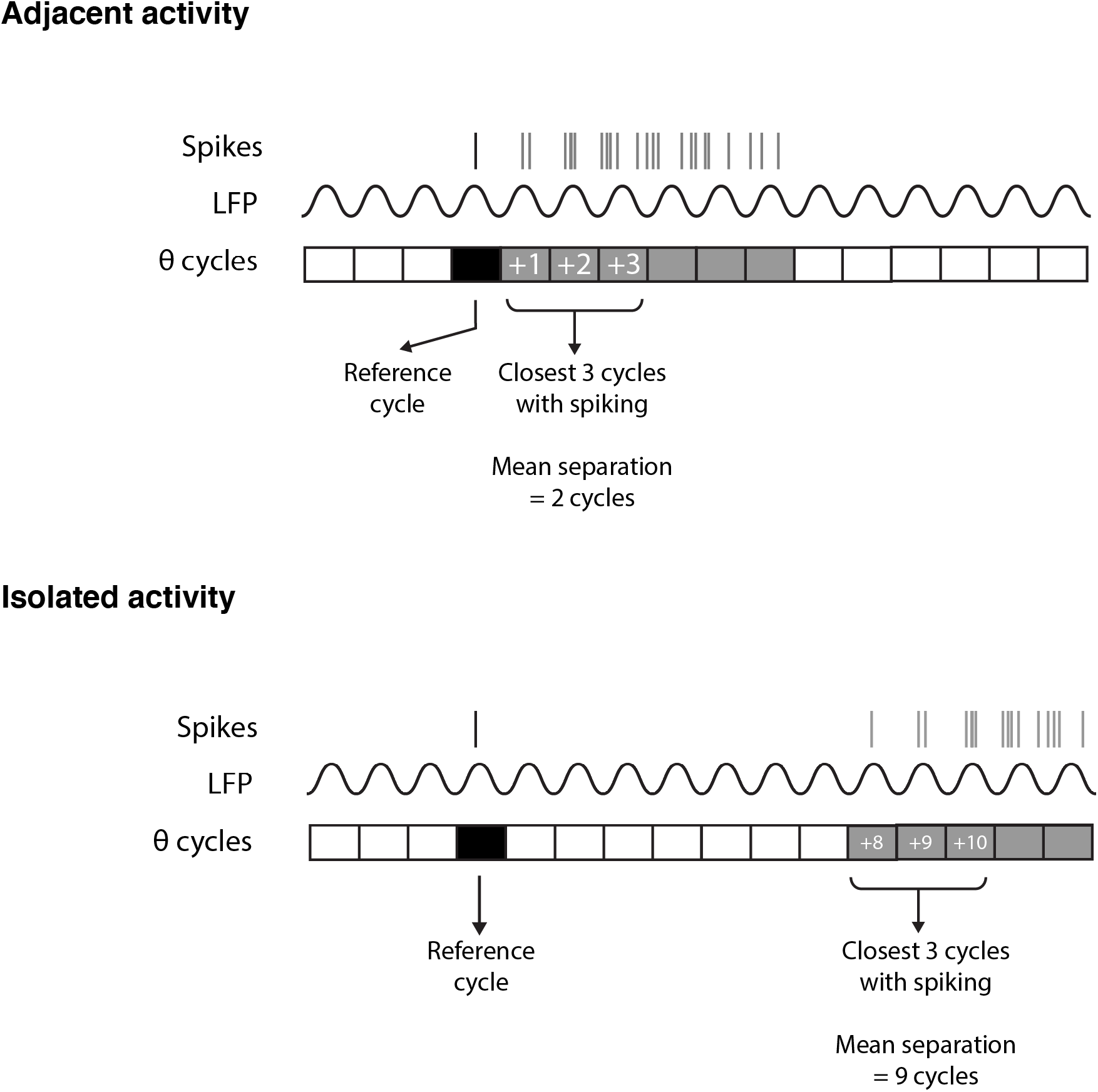
Adjacent and Isolated activity. Classification of adjacent versus isolated activity based on temporal separation between theta cycles with spiking.

**Figure S3.**
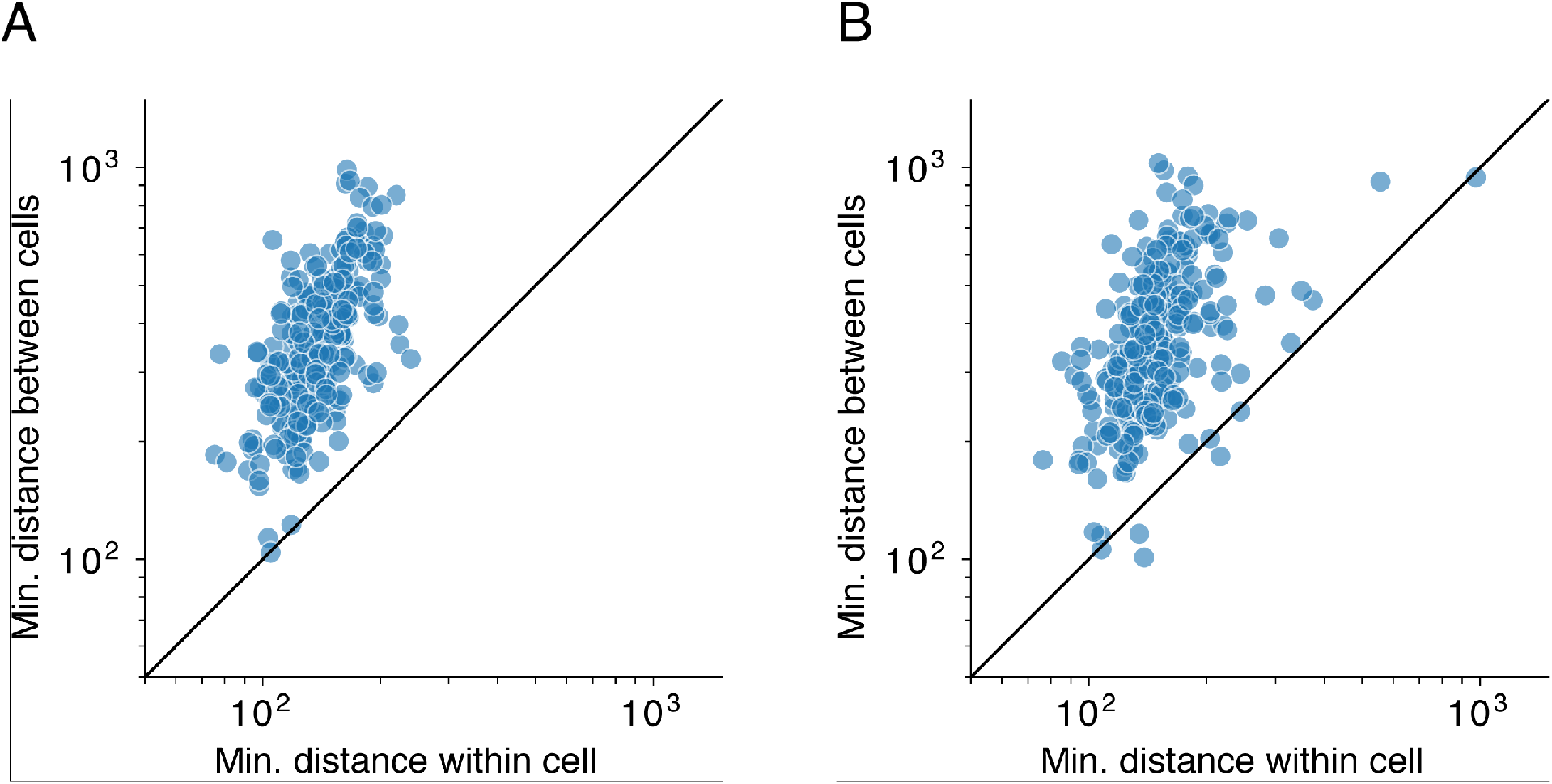
Isolated spiking activity is not due to spike assignment errors. A. Minimum Euclidean distance between the spike waveform of adjacent activity spikes within each cell versus all other place cell recorded on the same tetrode (n=260 individual cells compared to all other cells on the tetrode). Wilcoxon signed-rank test: p=2.9×10^−51^. B. Minimum Euclidean distance between the spike waveform of spikes classified as isolated activity within each cell versus all other place cell recorded on the same tetrode (n=260). Wilcoxon signed-rank test: p=8.8×10^−51^.

**Figure S4.**
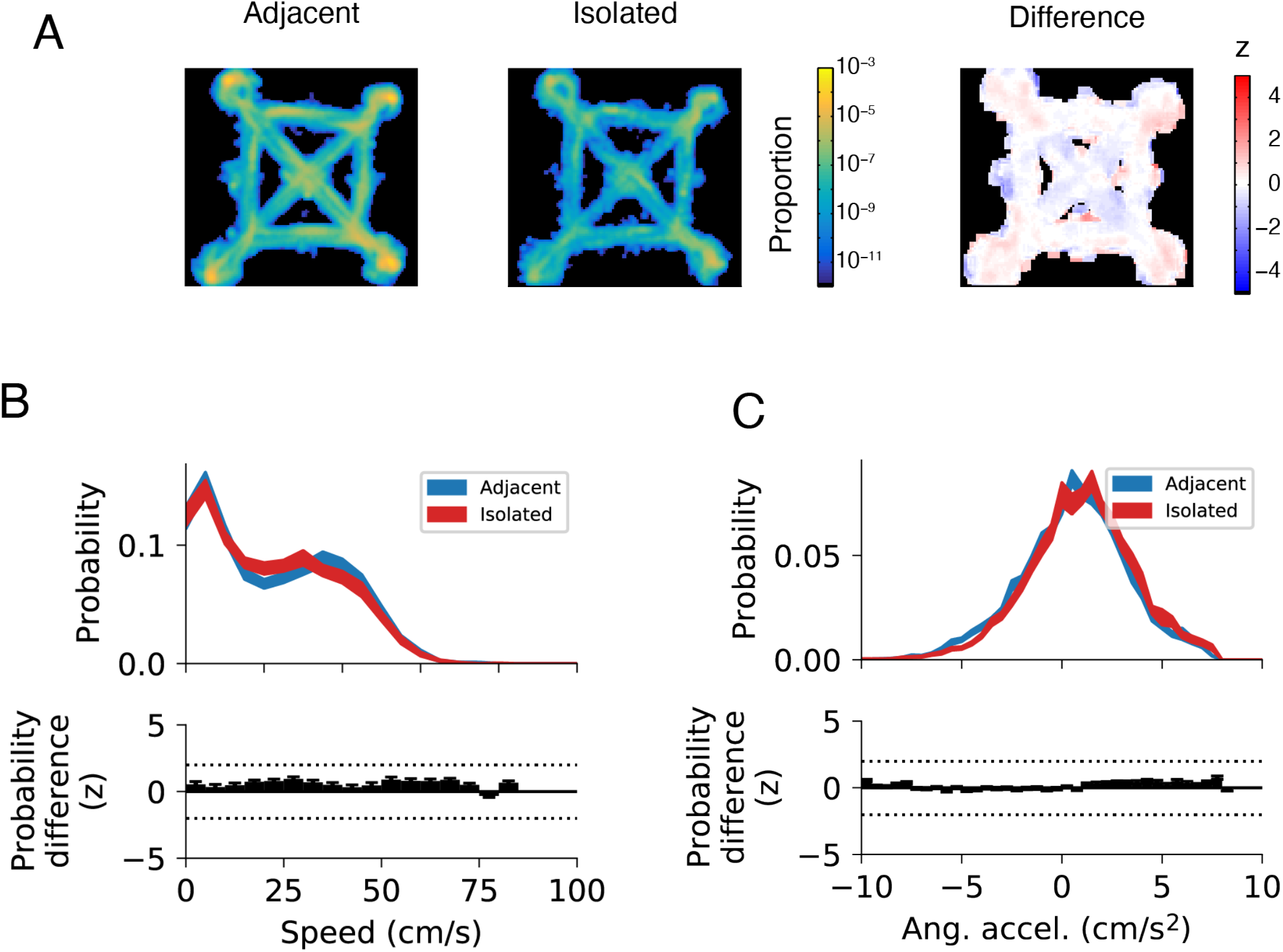
Animal location and movement correlates are similar for isolated and adjacent spiking. A. Normalized spatial distribution of theta cycles with adjacent (left) or isolated (center) spiking. Normalized difference between the spatial distributions (right). B. Distribution of animal speed (Mean ± SEM) at the time of adjacent or isolated activity (top). Significance of the difference (z) between the two distributions as determined using a permutation test (bottom). Dotted lines indicate ± 2z. C. Distribution of animal angular acceleration (Mean ± SEM) at the time of adjacent or isolated activity (top). Significance of the difference (z) between the two distributions as determined using a permutation test (bottom). Dotted lines indicate ± 2z.

**Figure S5.**
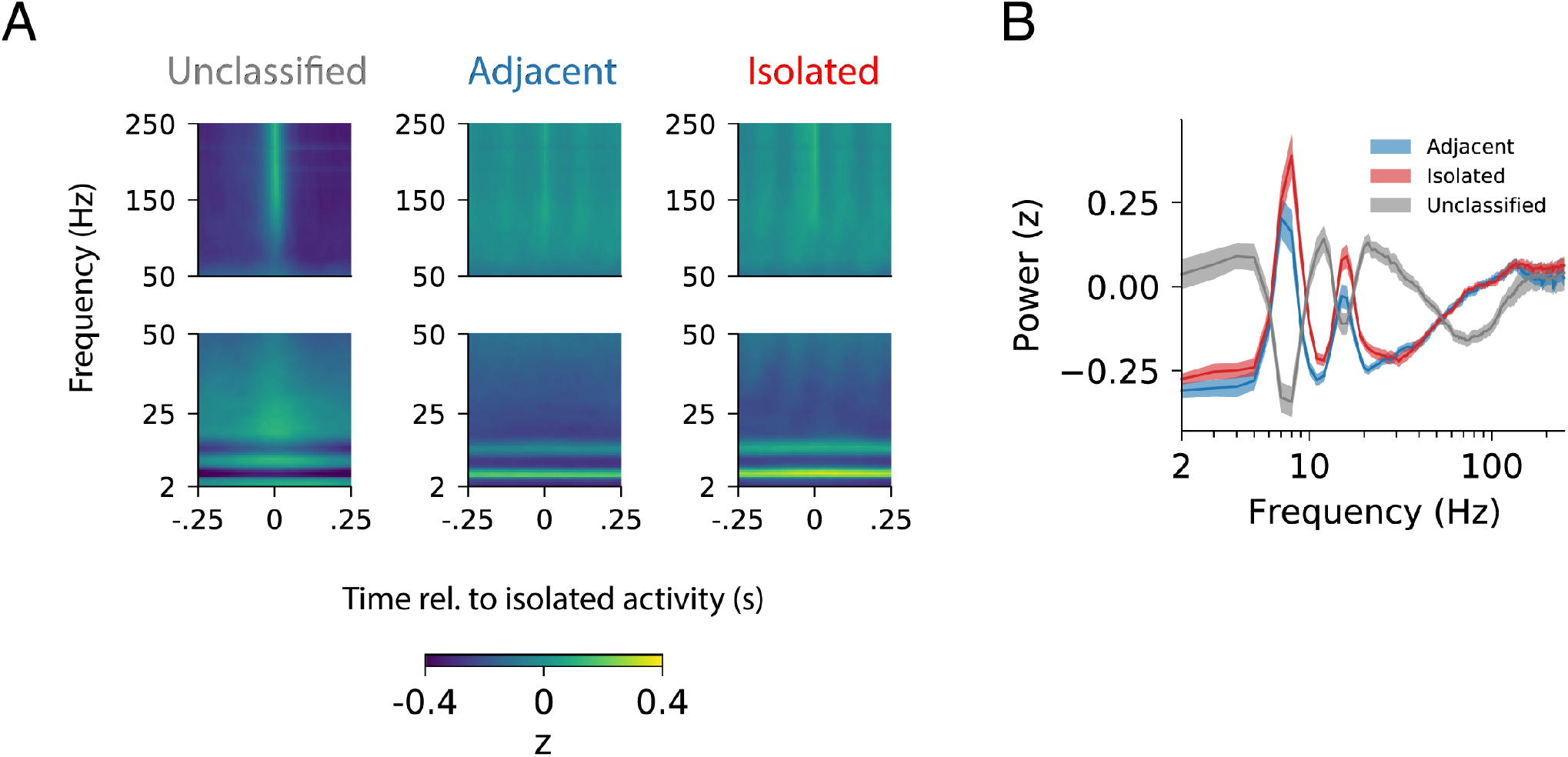
Network spectral signature around unclassified, adjacent and isolated spiking activity. A. Mean spike triggered spectrogram for unclassified (left), adjacent (center), and isolated (right) spiking activity (n=170 cells). Top panels show frequency ranges 50-250Hz. Bottom panels show frequency ranges 2-50Hz. B. Mean spectral power for a 50ms window centered at 0ms lag (Median ±95% CI).

**Figure S6.**
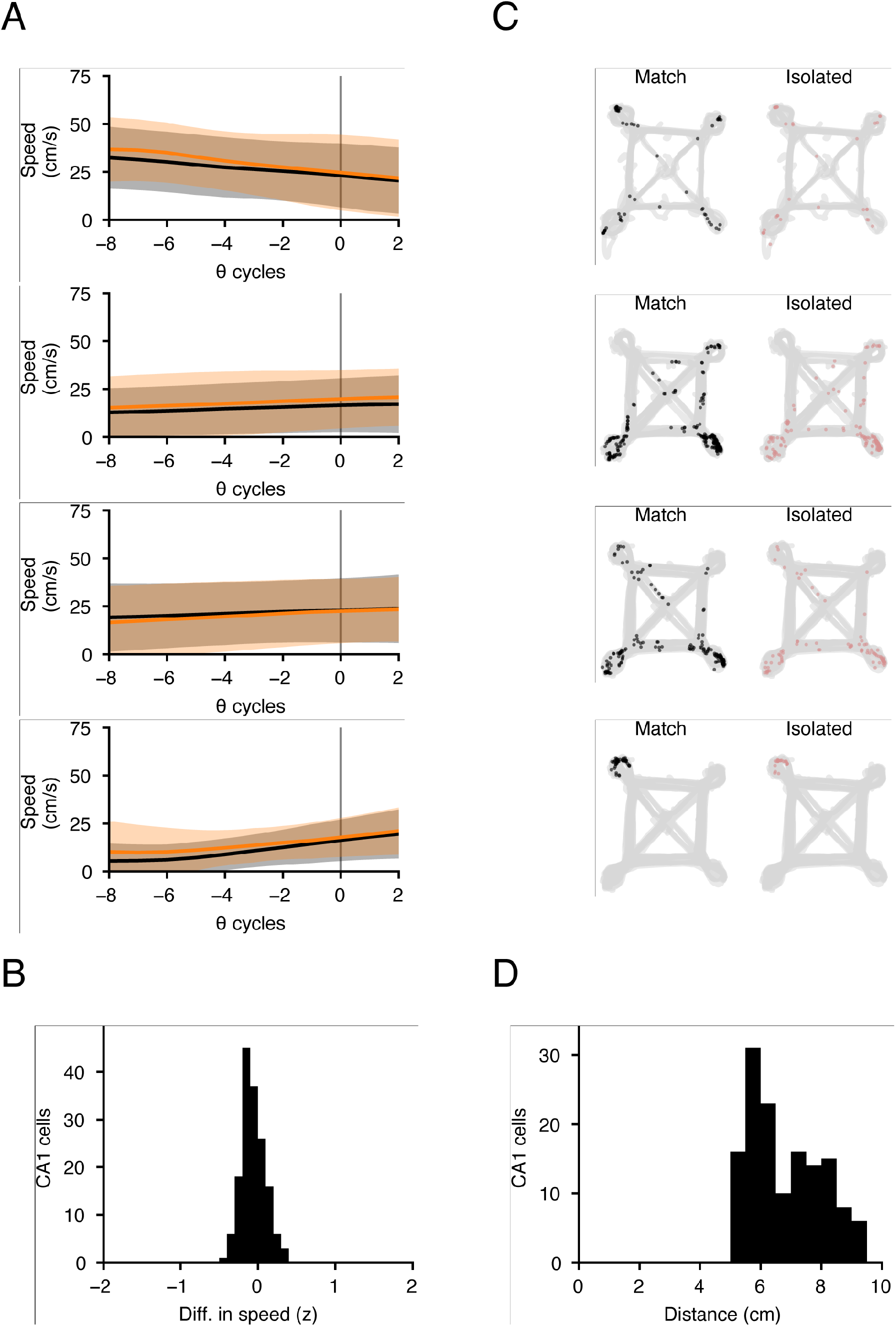
Each theta cycle with isolated place cell activity is matched with non-isolated cycles for speed, trajectory and location. A. Speed match profiles for examples in Figure 4B. B. Distribution of mean difference in speed between matched and isolated cycles. Speed profile of matched cycles were on average within −0.06 standard deviations of the speed profile of the isolated cycles. The difference is expressed as a z-score normalized against the speed distribution of isolated cycles. C. Location match profiles examples in Figure 4B. D. Distribution of the mean distance in cm between matched and isolated cycles. The location of the animal on matched cycles was on average 7.5cm from the location of the isolated cycle.

**Figure S7.**
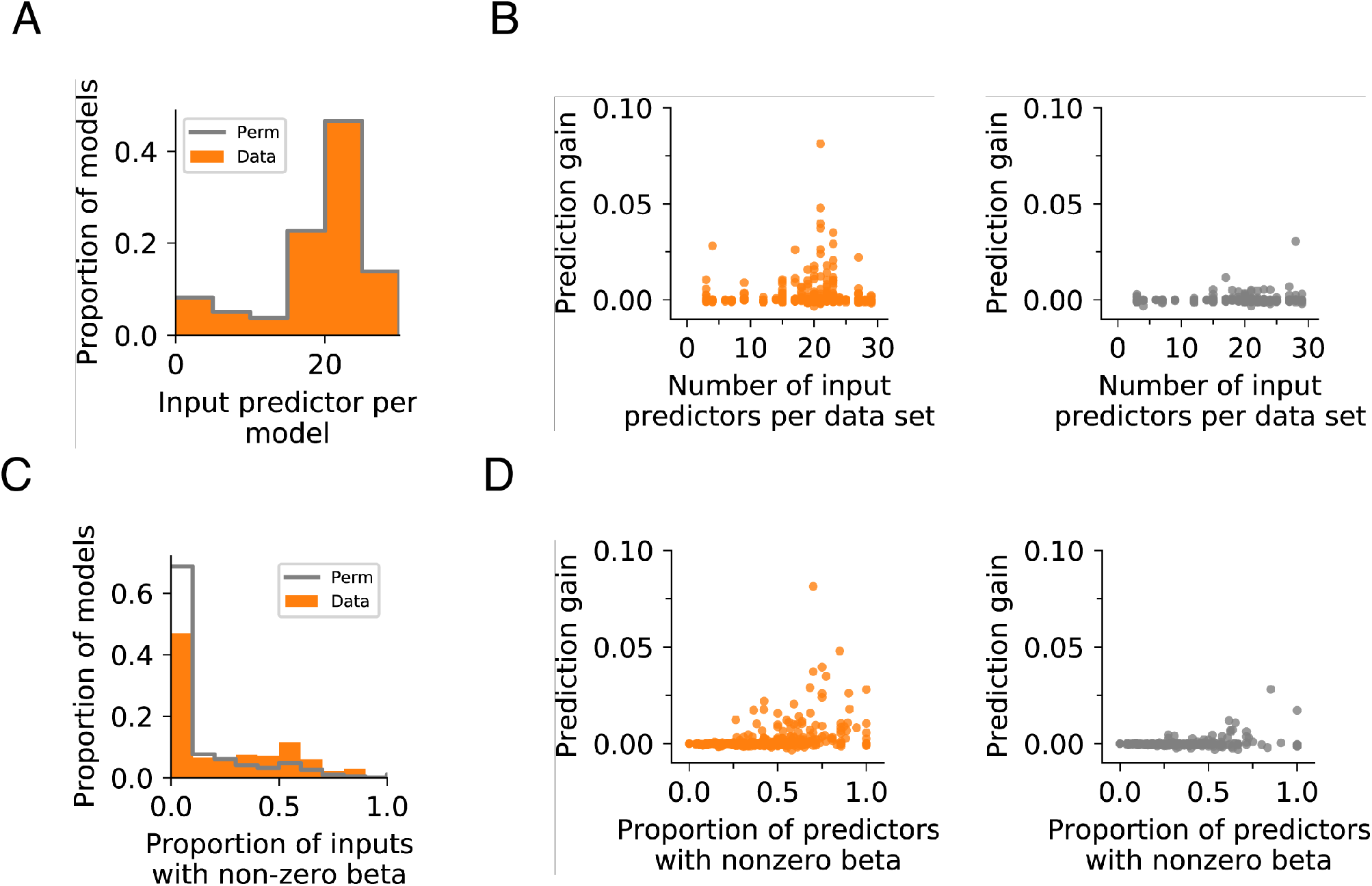
Model quality controls for GLMs using PFC activity to predict CA1 isolated activity. A. Input predictor count for actual and permuted datasets. Wilcoxon rank-sum test p=1.0. B. Prediction gain is not significantly correlated with the total number of input predictors used for prediction. Data: R^2^=0.00121, p=0.384; Permutation control: R^2^=0.00387, p=0.118. C. Models using actual data have higher proportions of predictors with non-zero β coefficients. Wilcoxon rank-sum test p=1.22×10^−16^. D. Prediction gain is positively correlated with the proportion of input predictors with non-zero beta coefficients. This is found in both actual (left) and permuted (right) datasets. Data: R^2^=0.133, p=3.10×10^−21^; Permutation control: R^2^=0.0537, p=0.3.80×10^−9^. Each point in the scatter represents a single fold of each model with 5 folds in total. All time points and models are shown.

## Methods

The data used in this study came from the same dataset used in previous publications (Yu, Kay et al. 2017, Yu, Liu et al. 2018).

### Animal and behavior

Experiments followed guidelines from the University of California San Francisco Institutional Animal Care and Use Committee and US National Institutes of Health. Six Long-Evans rats (male, 500-700g, 4-9 months of age) were first trained to traverse a linear track (1m) for reward (evaporated milk, Carnation brand, with 5% added sucrose). Next, the animals were trained on a foraging task ~21 days after surgery (Yu, Kay et al. 2017, Yu, Liu et al. 2018). Briefly, the task has four possible reward well locations, only two were chosen to deliver reward at a given time. The rat is trained to visit the two rewarded location in alternation to receive reward. The rewarded well locations changed within or between sessions, or between days. Two to three session (15-45 minutes) were performed each day with interleaved rest sessions (20-60 minutes). Reward was delivered (100-300μl at 20ml/min) using a syringe pump (NE-500 OEM, New Era Pump Systems Inc.) after the animal breaks an infrared beam at the well location.

### Implant

The recording drive was 3D printed (PolyJetHD Blue, Stratasys Ltd.) and contained up to 28 individually movable tetrodes. Tetrodes (Ni-Cr, California Fine Wire Company) were gold plated to 250kOhm at 1kHz.

The implanted recording drives targeted both dorsal CA1 (7 tetrodes) and dorsal PFC (14-21 tetrodes, housed in one cannula angled at 20 degrees toward the midline). CA1 AP: - 3.8mm and ML: 2.2mm. PFC (anterior cingulate cortex and dorsal prelimbic cortex): AP: +2.2mm, ML +1.5mm and DV between 1.88mm to 2.72mm depending on the AP and ML coordinates of each tetrode.

Initially, tetrodes were adjusted to reach the target DV coordinate (PFC) or guided by LFP and spiking patterns (CA1), every 2 days. Once the target was reached, tetrodes were adjusted (~30μm) to improve cell isolation at the end of an experiment day

### Histology

Recording sites were marked with electrolytic lesions by passing current through each tetrode (30μA, 3s) at the end of the experiment. Animals were perfused after 24 hours with paraformaldehyde (4% in PBS). The brain was removed, fixed (24 hours at room temperature), cryoprotected (30% sucrose in PBS at 4°C) and sectioned (coronal, 50μm). Cresyl Violet was used to stain the sections to identify sites of electrolytic lesions.

### Recording

Data were recorded with the NSpike system (LMF and J. MacArthur, Harvard Instrumentation Design Laboratory). Dim lighting was used during the experiment. An infrared LED array was mounted on the headstage amplifier to for position tracking. Video was recorded at 30Hz. We recorded LFP (0.5-400Hz at 1.5kHz) and spiking activity (600-6000Hz or 300-6000Hz at 30kHz) from each tetrode channel. For spike detection referencing, a tetrode located in corpus callosum was used for CA1 and a local tetrode without detected spikes was used for PFC.

### Data preprocessing

Manual spike clustering was performed based on peak amplitude, spike width and waveform principal components (MatClust, https://bitbucket.org/mkarlsso/matclust/src/master/).

To reconstruct the position of the animal, the centroid of the front and back diodes from the LED array was automatically extracted from the video.

### Spike clustering quality

To assess clustering quality, we analyzed the similarity in spike waveform within and across different units. We expect a well clustered unit to have spikes with waveforms that are similar to other spikes assigned to the same unit compared with spikes assigned to other units. Potential spike misassignment can occur only for spiking events detected on the same tetrode. We therefore compared spike waveforms from a given unit to spikes from all other units on the same tetrode. We computed the Euclidean distance between the spike waveforms (4 channels) for all pairs of spikes. Next, we compared the minimum waveform distance between spikes belonging to the same unit, and between that unit and all other units. This was done separately for spikes associated with isolated or adjacent theta cycles.

### Spatial spiking rate

The occupancy-normalized rate was calculated by dividing the number of spikes by the occupancy of the animal per spatial bin (2cm by 2cm) in the environment. A 2-dimensional symmetric Gaussian kernel (σ=2cm and 12cm spatial extent) was then used for smoothing.

### Theta cycle definition and classification

The theta-frequency component of the raw LFP signal was extracted using an equiripple finite impulse response band-pass filter (6-12Hz). Given theta is associated with movement states (Vanderwolf 1969), we used two criteria to exclude activity associated with immobility periods. First, we exclude periods when the speed of the animal was less than 2cm/s. Second, we excluded periods with Sharp Wave/Ripples, which occur during immobility or periods of slow movement, using previously described methods (Yu, Kay et al. 2017, Yu, Liu et al. 2018). For SWR detection, we used a speed threshold of <4cm/s to ensure SWRs occurring during intermediate speeds (>2cm/s but <4cm/s) are excluded. Spikes occurring during these excluded periods are classed as “Excluded”.

For spiking during the included periods, we classified each theta cycle and spikes belonging to that cycle as “isolated” or “adjacent” activity. This was done per place cell. The classification was based on the mean number of cycles separating a given cycle with spiking to its nearest 3 other neighboring cycles with spiking. A mean separation of 8 cycles is the threshold for classification as adjacent as opposed to isolated. This is based on the distribution of cycle separation across the entire place cell population.

### Theta cycle spatial distribution

To determine if isolated spiking occurred more frequently at certain locations in the environment compared with adjacent spiking, we first plotted the normalized spatial distribution of theta cycles containing each type of activity for each cell, averaged across the population. The spatial distribution of isolated and adjacent theta cycles for each place cell was calculated using the spatial spiking rate method described above, except with their respective cycles instead of spikes. To determine whether there are areas in the environment where the occurrence of adjacent and isolated activity differ, we applied a permutation technique. This involves first permuting the identity of each theta cycle labeled as having adjacent or isolated activity. The spatial distribution of the two permuted sets were calculated and subtracted from each other to obtain the difference. This was done 500 times to generate an expected distribution of differences. The actual difference in spatial distribution between isolated and adjacent activity was compared to the expected distribution and a z-score was calculated.

### Theta cycle movement correlates

To determine whether isolated and adjacent activity were associated with distinct movement correlates, we compared the distribution of animal speed and angular velocity at times of theta cycles containing each type of activity. This was done for each place cell and then averaged across the population. To determine if the two distributions were significantly different, we used a permutation approach. For each place cell, the identity of the theta cycle, whether it contains isolated or adjacent, was permuted. The difference between the two distributions permuted data sets was the recalculated. This was repeated 1000 times to obtain a distribution of expected differences. The actual difference was expressed as a z-score relative to the expected distribution.

### Theta phase locking analysis

The theta phase of each spike is relative to the phase of the reference signal obtained from the tetrode located in the corpus callosum. The mean phase preference for spiking activity for each place cell is the circular mean of the phases of all spike. Theta phase concentration is the magnitude of the vector sum of all spikes, where each spike is a unit vector with angle corresponding to the phase of theta.

### Time aligned spectrogram

To compute the spectral properties of network activity around spiking events we first used a bank of bandpass filters (center frequency ± 1Hz) to filter the LFP signal across the frequency range 2-250Hz. Each filtered signal normalized by subtracting the mean and dividing by the standard deviation. For each place cell, we then selected the normalized signal in a 500ms window centered on the time of each spiking event and averaged across spikes. This was repeated for spikes classified as belonging to isolated, adjacent theta cycles or excluded from analysis (see **Theta cycle definition and classification**). We ensured equal number of spikes were used to generate the average across spike types for each cell by sampling without replacement to match the type with the lowest count. The average for each place cell was then used to generate the mean for the entire population.

### Spiking coactivity

We quantified the likelihood a pair of place cells having isolated activity in the same theta cycle relative to the expected probability, similar to what has been done for SWRs (Cheng and Frank 2008, Singer and Frank 2009, Yu, Kay et al. 2017). The expected probability is the frequency of observing spiking from two cells in the same theta cycle given their relative frequency of spiking. For each cell, its spike count during a theta cycle with isolated activity was first binarized, where the cell was either spiking or not spiking in that theta cycle. The proportion of all theta cycles where both cells spiked was the observed coactivity. The expected coactivity was calculated by permuting the participation of each cell across all theta cycles with isolated activity. This was repeated 1000 times and to generate a distribution expected proportion of theta cycles with both cells have isolate activity. The observed proportion was converted to a z-score by subtracting the mean and dividing by the standard deviation of the expected distribution. This method accounts for the differences in the number of theta cycles with isolated activity for each cell in the pair.

To determine the temporal relationship between adjacent activity for a pair of place cells, we computed the cross-correlation between theta cycles with adjacent activity for a given cell pair. First, we assigned each theta cycle of a given cell as having adjacent spiking or not. Then, we cross-correlated the assignment for a pair of place cells, where the lag is measured in the number of cycles. We then found the absolute lag with the maximum cross-correlation value with for each place cell pair.

### Cycle matching

For each place cell, we matched each theta cycle with isolated activity with control theta cycles without spiking. These control cycles were drawn from other task trials from the same session and matched as closely as possible for trajectory, speed, and location. Two control cycles were selected for each actual cycle. Trajectory matching only included task trials where the animal performed the same trajectory and ensured the same direction of travel across all matched cycles. The speed matching process started with generating a reference speed profile distribution for a time interval around a theta cycle with isolated activity for a given cell. For each theta cycle with isolated activity, we then chose two candidate theta cycles without spiking. The speed profile for each candidate cycle around the same interval was compared with the reference distribution. The candidate cycle was accepted if the mean speed deviation compared with the reference distribution is <1σ The next inclusion criteria for the candidate cycle was having a location < 10cm from the theta cycle with isolated activity. This selection process was done without replacement. Only place cells with greater than 100 input cycles, including both isolated and matched cycles, were included in the analysis.

### Spiking normalization to theta cycles

For illustration purposes in Figure 4A, we converted PFC and CA1 spiking times to hippocampal theta cycle phases. Spiking times were transformed using linear interpolation from time to theta phase relative to the start of the theta cycle with isolated activity. The mean spiking rate was calculated with respect to theta cycles.

### Spiking rate comparison

We asked whether PFC spiking rate leading up to and including theta cycle with isolated compared with matched control cycles. For each CA1 cell, we first identified cycles with isolated spiking and control cycles without isolated spiking (see above). We next found all PFC cells that were simultaneously recoded with the CA1 cell. For each of these PFC cells, we compared the spike count in time intervals leading up to the theta cycle with or without isolated spiking from the CA1 cell. Under the null hypothesis, the difference between the two sets of spike counts will not be significantly different than chance. To estimate the significance of the spike count difference, we used a permutation test where we permuted the theta cycle identity 1000 times and calculated the difference between the PFC spiking for each permutation. The actual difference was expressed as a z-score relative to this permuted distribution by subtracting the mean and standard deviation of the permuted distribution. As an additional control to estimate the expected difference between the groups, we repeated the analysis by first generating a permuted data set where the theta cycle identity (with or without isolated spiking) was permuted. This difference in the spiking rate of this permuted data set, expressed as a z-score, was calculated as the actual data set.

### Generalized Linear Models

We asked whether spiking activity from simultaneously recorded PFC cells can predict the occurrence of isolated activity from a CA1 cell. We built cross-validated Generalized Linear Models (GLMs) (Binomial distribution with logit link function) with elastic-net regularization, which combined LASSO and Ridge regularization to reduce overfitting (Zou and Hastie 2005). To do this, we first identified theta cycles with isolate spiking for a CA1 cell. We next identified another control set of theta cycles when the CA1 cell did not spike. These control cycles were matched for animal speed, movement direction and location (see **Cycle matching**). We created a model for each CA1 cell to determine whether PFC spiking activity can distinguish between cycles with or without isolated spiking in a time window relative to the cycle with isolated spiking. We first modelled using activity in the 12 cycles previous to the cycle with isolated activity and then the 12 cycles after the cycle with isolated activity. We ensured no other isolated activity occurred in this window used for prediction. A 4-cycle bin size was used for grouping PFC activity since PFC activity shows relatively long autocorrelation times.

### Modelling parameters

MATLAB’s *lassoglm* function was used (‘distr’=‘binomial’, ‘Link’=‘logit’). The optimization was equally weighed between LASSO and Ridge methods (‘alpha’=0.5). Shrinkage parameter (λ) optimization was done using 3-fold cross-validation (‘CV’=3) with 5 Monte Carlo repetitions (‘MCReps’=5). We used 5-fold cross validation and averaged the outcome across the 5 cross-validations.

### Prediction gain

Prediction gain describes the whether the models can predict the outcome above chance. For the actual dataset (Prediction gain_Data_), we did this by first calculating the mean absolute error (MAE) between the predicted and actual outcome for the validation partition (MAE_Data_). To estimate chance performance, we repeated the prediction 5000 times, each time with the outcome permuted, and calculated the MAE. The chance MAE is the mean MAE of the 5000 control predictions (MAE_Permuted_). The prediction gain is log_10_(MAE_Permuted_/MAE_Data_). A positive prediction gain means the Data group had a smaller error, or better prediction, compared with the Permutated group (Rothschild, Eban et al. 2017).

We also used a second approach to estimate chance prediction. Instead of building the model using actual data, we permuted the trial identity of PFC input, which preserves the input spiking distribution but destroys any potential relationships between trials and the outcome in CA1. We repeated the entire modelling procedure using permuted data and calculated the prediction gain (Prediction gain_Permuted_).

To estimate whether there is significant above chance prediction of CA1 isolated activity from PFC activity, we performed a permutation test (n=10000 permutations) on the mean prediction gain between the actual (Prediction gain_Data_) and permuted (Prediction gain_Permuted_) datasets. The Bonferroni correction was used to adjust the significance of the prediction to account for multiple comparisons between time windows.

### PFC predictive ensemble correlation

To determine whether there is specificity in the coordination between PFC and CA1 around the time of isolated activity, we asked if isolated activity for each CA1 cell was predicted by activity from distinct combination of PFC cells. This was done by calculating the Pearson’s correlation between the β coefficients of CA1 models that were generated from data recorded on the same day. We selected models that yielded a prediction gain >1, had a minimum of 5 predictors and had at least 2 CA1 models from the same day. This produced a dataset from 17 days, with a median of 3 (Q1: 2, Q3: 5) models per day, with 21 (Q1: 19, Q3: 23) predictors per model, and with a median prediction gain of 1.014 (Q1: 1.0015, Q3: 1.026).

### Model quality assessment

We checked the quality of our models by examining the relationship between prediction gain and the contribution of predictors to the model. For these linear models, we used the value of the β coefficient to indicate the contribution of a predictor to the prediction, where predictors with non-zero β coefficient may contribute to the prediction. We examined how the prediction value varied with the proportion of predictors with non-zero β coefficients, or the total number of input predictors, using linear regression. We also compared these relationships between models with actual data or permuted data. Models with greater predictive power are expected to have higher proportion of predictive features whereas the number of input predictors should not affect the outcome.

### Statistical Analyses

Circular statistical analyses were performed using the Circular Statistics Toolbox in MATLAB (Berens 2009). Statistical tests were performed using standard MATLAB modules and Scipy Statistical Functions (scipy.stats). All tests were two-sided.

